# Unsupervised learning for robust working memory

**DOI:** 10.1101/2021.05.17.444447

**Authors:** Jintao Gu, Sukbin Lim

**Affiliations:** Neural Science, New York University Shanghai, 1555 Century Avenue, Shanghai, 200122, China; NYU-ECNU Institute of Brain and Cognitive Science at NYU Shanghai, 3663 Zhongshan Road North, Shanghai, 200062, China

## Abstract

Working memory is a core component of critical cognitive functions such as planning and decision-making. Persistent activity that lasts long after the stimulus offset has been considered a neural substrate for working memory. Attractor dynamics based on network interactions can successfully reproduce such persistent activity. However, it suffers from a fine-tuning of network connectivity, in particular, to form continuous attractors suggested for working memory encoding analog signals. Here, we investigate whether a specific form of synaptic plasticity rules can mitigate such tuning problems in two representative working memory models, namely, rate-coded and location-coded persistent activity. We consider two prominent types of plasticity rules, differential plasticity targeting the slip of instant neural activity and homeostatic plasticity regularizing the long-term average of activity, both of which have been proposed to fine-tune the weights in an unsupervised manner. Consistent with the findings of previous works, differential plasticity alone was enough to recover a graded-level persistent activity with less sensitivity to learning parameters. However, for the maintenance of spatially structured persistent activity, differential plasticity could recover persistent activity, but its pattern can be irregular for different stimulus locations. On the other hand, homeostatic plasticity shows a robust recovery of smooth spatial patterns under particular types of synaptic perturbations, such as perturbations in incoming synapses onto the entire or local populations, while it was not effective against perturbations in outgoing synapses from local populations. Instead, combining it with differential plasticity recovers location-coded persistent activity for a broader range of perturbations, suggesting compensation between two plasticity rules.

**Author Summary:** While external error and reward signals are essential for supervised and reinforcement learning, they are not always available. For example, when an animal holds a piece of information in mind for a short delay period in the absence of the original stimulus, it cannot generate an error signal by comparing its memory representation with the stimulus. Thus, it might be helpful to utilize an internal signal to guide learning. Here, we investigate the role of unsupervised learning for working memory maintenance, which acts during the delay period without external inputs. We consider two prominent classes of learning rules, namely, differential plasticity, which targets the slip of instant neural activity, and homeostatic plasticity, which regularizes the long-term average of activity. The two learning rules have been proposed to fine-tune the synaptic weights without external teaching signals. Here, by comparing their performance under various types of network perturbations, we reveal the conditions under which each rule can be effective and suggest possible synergy between them.

## Introduction

Continuous attractors have been hypothesized to support brains’ temporary storage and integration of analog information (1–3). An attractor is an idealized stable firing pattern that persists in the absence of stimuli, and integration is allowed if these attractors form a continuous manifold. Theoretical models predict that neural activity should be restricted within but free to move along this manifold, making stochastic fluctuation correlated among neurons, as is validated in the brainstem oculomotor neural integrator (4), the entorhinal grid cell system (5), and prefrontal visuospatial selective neurons (6).

Computationally, the performance of continuous attractors is known to be sensitive to network parameters, which is termed as the “fine-tuning problem” (7,8). The slight imperfection of synaptic weight asymmetry could make continuous attractors break down into a few discrete attractors or cause an overall drift of activities. This raises the question of how continuous attractors could exist in the brain. Noting that the model is just an idealization, earlier studies have proposed that continuous attractors can be approximated by finely discretized attractors with a hysteresis of coupled bi-stable units, which would make the system more robust (9,10). Recent theoretical studies suggest other complementary mechanisms, including derivative feedback and short-term facilitation, with the former slowing down activity decay (11,12) and the latter transiently enhancing stability (13,14).

These workarounds could make continuous attractors more tolerant to parameter perturbation. Not mutually exclusively, long-term plasticity is believed to take part in settling a reasonable parameter range. For example, the plasticity involved in the fish oculomotor integrator has been most studied. Previous works have proposed either visually supervised plasticity (15–17) or self-monitoring plasticity acting in the dark (18,19). These plasticity rules utilize time-derivative signals to detect slip of eye position or neural activity. Note that similar mechanisms can be generalized to mediate the tuning conditions of the parametric working memory encoding analog information (11,17,20). More broadly, derivative-based rules have been suggested to learn temporal relationships between input and output (21–23) and in reinforcement learning (24–26).

Another class of long-term synaptic plasticity suggested for continuous attractors is homeostatic plasticity, which regularizes the excitability of neurons (27). Many models focused on the role of homeostatic plasticity to prevent instability. As homeostatic plasticity tends to pull excitation down or boost inhibition when network activity is higher than a reference value, the positive feedback between network activity and activity-dependent plasticity can be counterbalanced (28). On the other hand, Renart et al. (29) considered network storing spatial information in a spatially localized “bump” activity pattern and proposed an additional role of homeostatic plasticity, that is to regularize the network patterns and recover tuning condition for spatial working memory perturbed by the heterogeneity of local excitability. Similarly, Pool and Mato (30) suggested that for developing orientation selectivity through Hebbian learning in recurrent connections, homeostatic plasticity can enforce symmetry in synaptic connections such that all orientation can be represented equally in the networks.

Both differential and homeostatic plasticity suggested for attractor networks are unsupervised. External supervisory or reward signals are not required to recover the required tuning condition. As shown previously, they can act after the offset of sensory signals and might be suitable for memory tasks that typically have a long memory period without external input. However, previous works have investigated the effect of differential plasticity and homeostatic plasticity partially for different types of continuous attractor or under particular types of perturbations in the network parameters.

Therefore, we investigated whether these two forms of learning can recover persistent activity in continuous attractors, which require fine-tuning conditions of network parameters. As a systematic study, we considered two different types of continuous attractors, namely, rate-coded and location-coded persistent memory, in a single framework, called the negative derivative feedback mechanisms (11,12). First, we formally described the fine-tuning problem in a rate-coded attractor system with a simpler network architecture than a location-coded attractor. We examined the effects of differential plasticity and homeostatic plasticity and how recovery from perturbation in connectivity depends on the learning parameters. Then we extended the scope of our investigation to a location-coded system that requires spatially structured networks and investigated the recovery of tuning conditions under various types of perturbations. Finally, we demonstrated that two rules could partially compensate for each other when they are combined.

## Results

### Rate-coded persistent activity in one homogenous population

Before we discuss the synaptic plasticity rule that stabilizes persistent spatial patterns of activity, we first consider the similar mechanism applied for a rate-coded persistent activity where the persistent firing rate of memory neurons varies monotonically with the encoded signals (2). Compared to location-coded memory suggested for maintaining spatial information, the rate coded one has been suggested to maintain a graded level of information such as somatosensory vibration frequency (31,32). Previous theoretical works proposed that recurrent circuits can maintain both types of memory based on similar feedback mechanisms despite the different network architecture (12). Thus, we first gain insight into how the specific form of synaptic plasticity can stabilize persistent memory in the rate coding scheme, which has a simpler network structure.

As the rate-coded network can be built upon a spatially homogeneous structure, its dynamic principle can be captured in the mean-field equations describing the network dynamics with one variable (see Methods). Two representative feedback mechanisms have been proposed based on recurrent network interactions, positive feedback, and negative derivative feedback, both of which is described by the following equation,

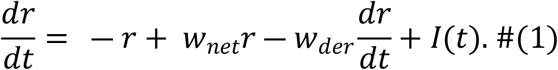

In the above equation, *r* represents the mean firing rate of the network activity. We considered time *t* and other time constants are unitless (normalized with the intrinsic time constant of *r*) for simplicity. The first and last terms on the right side represent the intrinsic leakage and transient external input. The second and third terms represent the feedback arising from recurrent inputs.

In the positive feedback models, the excessive excitatory inputs need to be tuned to cancel the intrinsic leakage such that the net gain w_net_ in the second term is tuned to be one, whereas w_der_ is typically zero (11). On the other hand, in the negative derivative feedback models, balanced excitatory and inhibitory recurrent inputs with different kinetics generate the resistive force against memory slippage similar to time-derivative activity in the third term. As its strength represented by w_der_ increases while the second term remains relatively small, the effective time constant of decay of network activity increases proportionally. Thus, for large negative derivative feedback, the decay of activity slows down (11).

With a long time-constant of decay, both networks show integrator-like properties such that during the stimulus presentation, it integrates the external input. After its offset, it maintains persistent activity at different levels (Fig. 1A). However, any memory circuits keeping the information in continuum states face a fine-tuning problem (7,8,33). Similarly, for rate-coded persistent memory, despite the different tuning conditions in positive feedback models and negative-derivative feedback models, the deviation from the perfect tuning leads to a gross disruption of persistent activity. For instance, a reduction in the E-to-E connection mimics the effect of NMDA perturbation in memory cells shown experimentally (11,34). Such a perturbation causes an imbalance between the recurrent excitation and inhibition in negative derivative feedback models and leads to the rapid decay of the activity (Fig. 1B).

**Figure 1.**
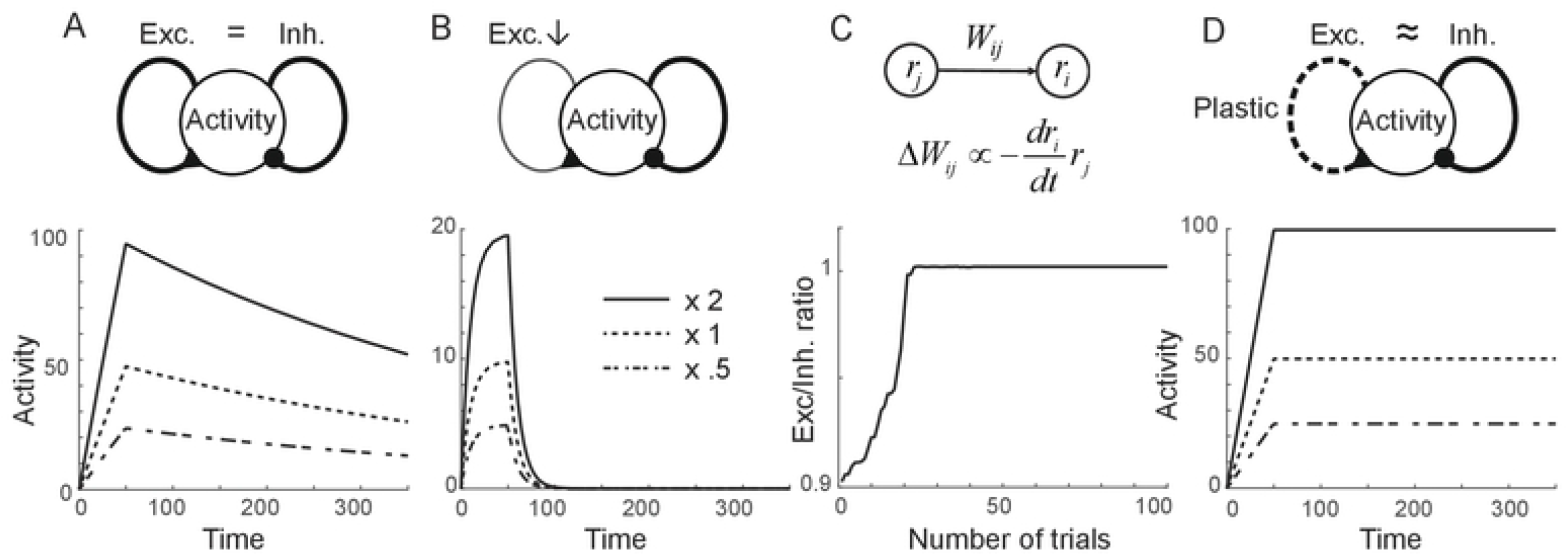
Recovery of rate-coded persistent activity through differential plasticity. A: Maintenance of persistent activity through negative-derivative feedback. With balanced excitation and inhibition with slower excitation, the network can maintain persistent activity at different rates (solid, dotted, and dash-dotted curves represent activity for different input strengths). B: Disruption of persistent activity (bottom) under the perturbation in the recurrent excitatory connections (arrow in top panel). C: Schematics of differential plasticity (top) and recovery of E-I balance under differential plasticity (bottom). D: Maintenance of persistent activity after the recovery of E-I balance through differential plasticity.

### Stabilization of persistence through differential plasticity

To mitigate this fine-tuning condition and to make the network resilient against perturbations, several forms of synaptic plasticity have been proposed. Two prominent synaptic plasticities suggested for persistent activities are homeostatic plasticity (27,29) and differential plasticity (18,19). Here, we examine how each plasticity can stabilize a rate-coded persistent activity.

First, we consider differential synaptic plasticity where the synaptic update depends on the firing rate and its time derivative of pre- and postsynaptic activities [(18); Fig. 1C]. Previous work showed that such a plasticity rule updates the synaptic connection to reduce the overall derivative of network activities (18). We considered the negative-derivative feedback model composed of one homogenous population to understand further how the fine-tuning condition can be achieved through the differential plasticity rule. We assumed that the network receives balanced recurrent excitatory and inhibitory inputs with its strengths denoted as *W*_*exc*_ and *W*_*inh*_, and excitatory inputs have slow kinetics than the inhibitory inputs. If initially balanced excitation and inhibition is perturbed by the reduction in the excitatory connection and excitatory connection changes according to the differential plasticity rule, the dynamics of the system can be captured by the firing rate *r* and excitatory connection strength *W*_*exc*_ as

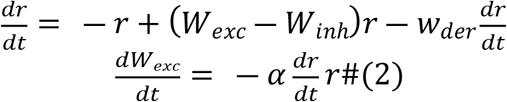

where *w*_*der*_ is proportional to *W*_*inh*_ and the difference of the time constants for excitatory and inhibitory inputs feedback (Methods).

The steady states of the system are *r* = 0 or *dr*/*dt* = 0, where the latter can be achieved for balanced excitation and inhibition, that is, *W*_*exc*_ becomes closer to *W*_*inh*_ for large *W*_*inh*_. However, in successive trials where each trial is composed of stimulus presentation and delay period, and assuming that external input during the stimulus presentation reset *r* without changing *W*_*exc*_, we found that only *dr*/*dt* = 0, that is, the balanced tuning condition can be achieved (Fig. 1C,D). Once this tuning condition is achieved, the network can maintain the graded level of persistent activities (Fig. 1D).

We further investigated how the recovery of a tuning condition depends on the parameters of synaptic plasticity by examining the phase plane of *r* and *W*_*exc*_ (Fig. 2A). The learning speed α and *W*_*inh*_ have similar effects of modulating the vector field along the *W*_*exc*_-axis such that increasing *W*_*inh*_ is effectively the same as decreasing α (Fig. 2B-C; Methods). In other words, stronger derivative feedback requires a longer time to recover after the same percentage of perturbation, so the recovery duration should scale with synaptic strength *W*_*inh*_. On the other hand, larger perturbation leads to the initial *W*_*exc*_ further away from the balanced state, making it longer to recover its tuning condition (Fig. 2D). Finally, overall input strengths during the stimulus presentation determine the magnitude of *r* to be reset during the stimulus presentation such that larger stimulus strength pushes the system in a faster speed regime and makes the system faster to converge (Fig. 2E).

**Figure 2.**
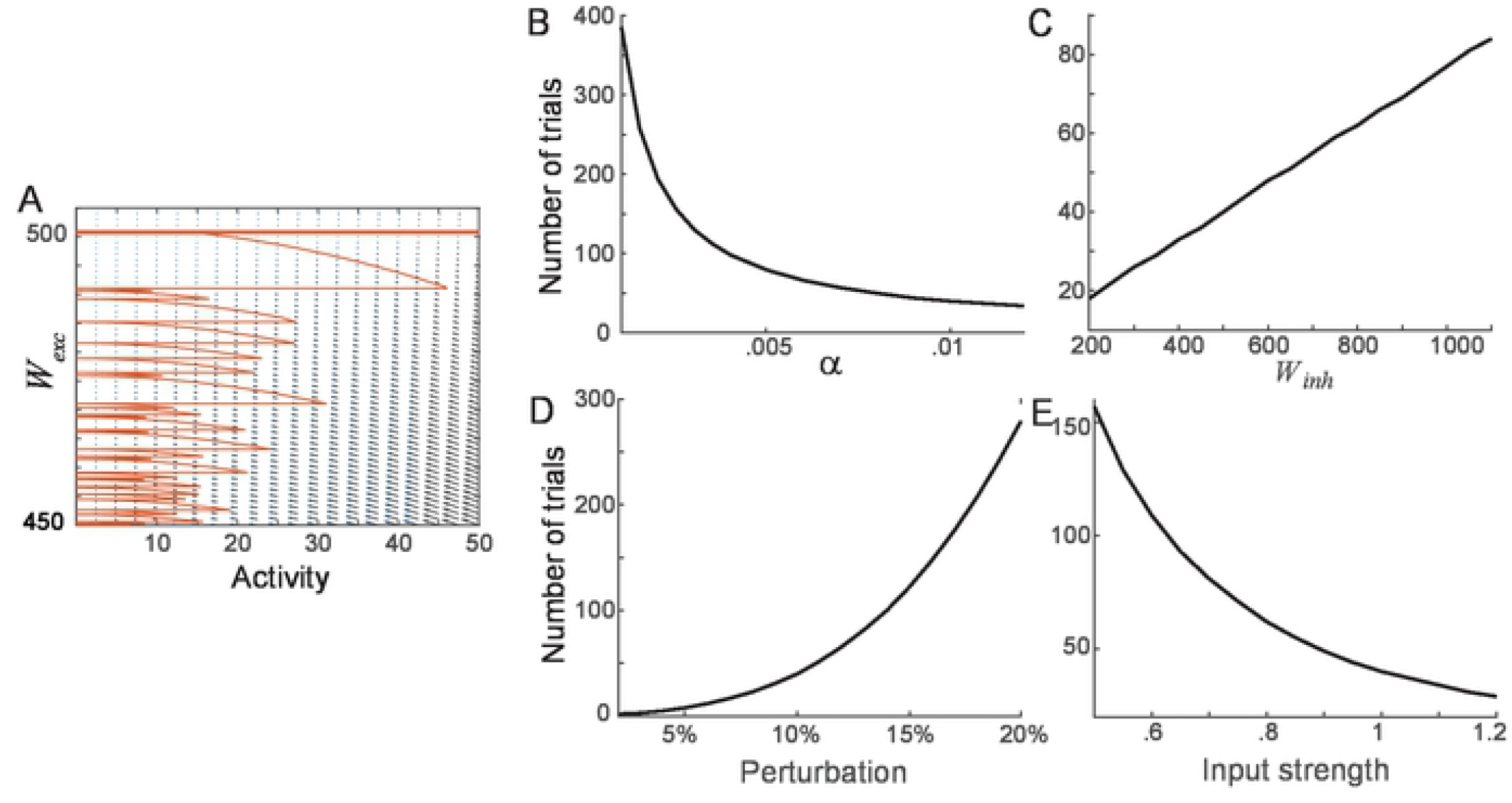
Recovery dynamics dependence on learning parameters under differential plasticity. A: Phase-plane of activity *r* and synaptic strength of recurrent excitation *W*_*exc*_. The black arrows represent a vector field for the dynamics of *r* and *W*_*exc*_, described in Eq. 2. The red curve is a trajectory starting from 10% perturbation in *W*_*exc*_, that is, *W*_*exc*_ = 0.9*W*_*inh*_ with *W*_*inh*_ = 500. During the stimulus presentation, the trajectory jumps horizontally, and input strengths vary randomly across trials. B-D: Dependence of recovery speed on learning and network parameters. The minimum number of trials for *W*_*exc*_ to reach up to 1% precision was obtained by varying the learning speed α (B), *W*_*inh*_ (C), perturbation strength (D), and relative mean input strengths across the trial (E).

### Homeostatic plasticity is effective but sensitive

While differential plasticity has been shown to stabilize the rate-coded persistent activity (11,18,19), homeostatic plasticity has been suggested to stabilize different forms of memory, such as spatial working memory (29) and discrete working memory (35,36). The homeostatic plasticity regulates the excitability of postsynaptic neurons such that in its typical form, all incoming synapses onto the postsynaptic neurons multiplicatively scale for the long-term average rate to achieve their target firing rates *r*_0_ (Fig. 3A). As for differential plasticity, we examined the effect of homeostatic plasticity in one homogenous population for a rate-coded persistent activity, whose dynamics is described as

**Figure 3.**
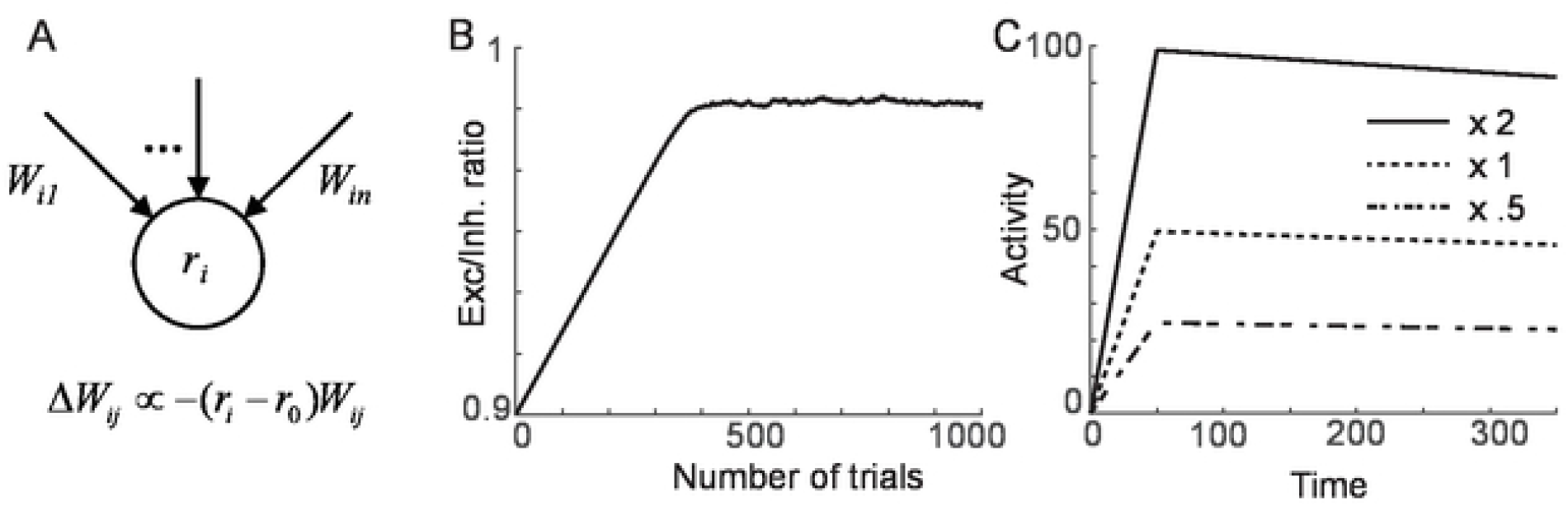
Recovery of rate-coded persistent activity through homeostatic plasticity. A: Schematics of homeostatic plasticity scaling the strengths of incoming synapses to achieve the target firing rate *r*_*0*_. B-C: Recovery of E-I balance (B) and maintenance of persistent activity at the different levels after the recovery (C).

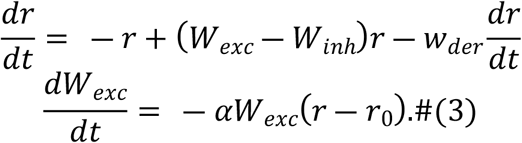

The steady-state of such a system is achieved when *r=r*_*0*_ and *dr*/*dt* = 0, that is, *W*_*exc*_ ≈ *W*_*inh*_ for large *W*_*inh*_. Note that this is more stringent than those for differential plasticity that requires the latter balance condition only.

Similarly to differential plasticity, we found that the steady-state can be achieved through homeostatic plasticity. However, it requires additional tuning of input strengths. The input strength set the initial condition for *r* at the beginning of the delay and the mean of initial *r* needs to be *r*_*0*_ on average. In such a case, the network achieves the balance condition (Fig. 3B) and maintains the rate-coded persistent activity (Fig. 3C). However, for inadequately tuned input, the steady-state cannot be achieved (Fig. 4A-B). For the mean of the input strength making *r* in the beginning of the delay period smaller (larger) than *r*_0_, the dynamics of *r* drifts upward (downward) to achieve *r*_0_ on average during the delay period (Fig. 4A-B, lower panels). Consequently, *W*_*exc*_ is stabilized to be excessive (deficient) compared to *W*_*inh*_ (Fig. 4A-B, upper panels). Thus, the homeostatic rule for rate-coded persistent memory requires tuning of input strengths and duration to achieve its target rate *r*_0_ and balance condition of recurrent excitation and inhibition.

**Figure 4.**
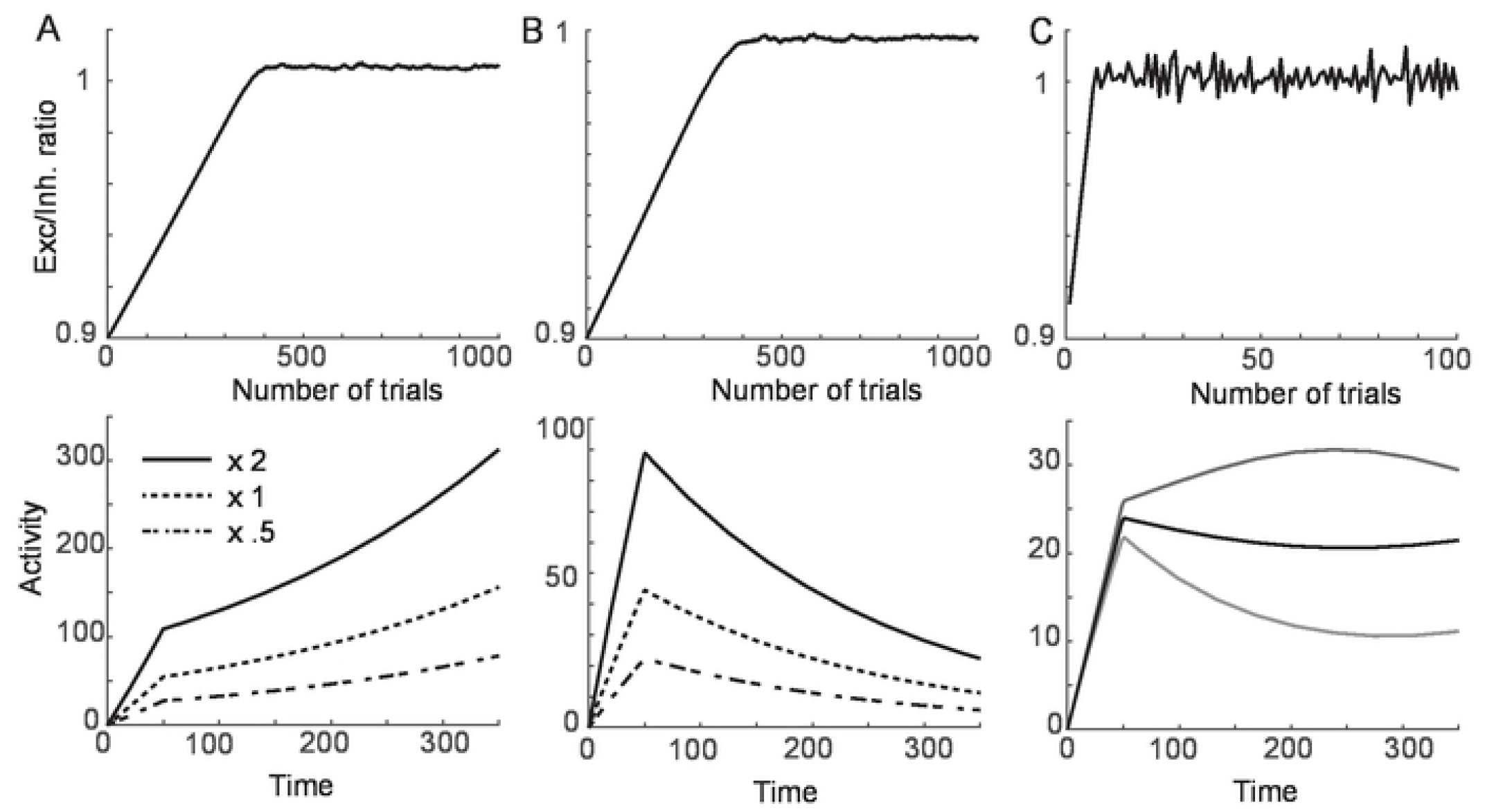
Sensitivity of homeostatic learning rule on learning parameters. A-B: Sensitivity to mismatch between *r*_0_ and input strengths. With lower mean input strengths compared to those in Fig. 3, the mean firing rate at the beginning of the delay period is lower than the target firing rate *r*_*0*_, and the dynamics drift upward to achieve *r*_*0*_ on average during the delay period (A, bottom). This results in excessive *W*_*exc*_ compared to *W*_*inh*_ (A, top). The opposite leads to the decay of activity and deficient *W*_*exc*_ (B). C: Sensitivity to the learning rate. For a faster learning rate, the homeostatic plasticity leads to the oscillation even for property tuned inputs, and the activity can vary across different trials for the same strength of the input.

Also, it is notable that the stability analysis further reveals that near the steady-state, the system shows damped oscillation. Its frequency depends on the speed of the homeostatic learning rule such that faster learning leads to faster oscillation. In successive trials with reset in *r*, the faster learning leads to the ongoing oscillation near the steady-state even for a properly tuned input such that for different trials, the dynamics cannot be stabilized (Fig. 4C). Overall, the analysis of one homogenous population shows that although homeostatic rule can recover persistent activity for rate-coded memory, it is sensitive to input parameters and learning speed.

### Location-coded persistent memory in spatially structured network

So far, we showed how two prominent plasticity rules could stabilize rate-coded persistent memory in one homogenous population. However, whether the same mechanism can be generalized to stabilize location-coded persistent memory is in question because both rules are local, depending on pre- and postsynaptic activity but have no regularization on a spatial pattern of activities required for encoding spatial information. Here, we considered the negative derivative feedback model suggested for spatial working memory (12) and explored under which condition such generalization can be made.

Previous work showed that the principle for negative derivative feedback found for one homogenous population could be extended to the network with a functionally columnar structure required to maintain a spatial pattern of persistent activity. Consistent with experimental observations (37–39), both excitatory and inhibitory neurons in each column have similar spatial selectivity. The connectivity strengths decrease as the preferred features over the columns get dissimilar (Fig. 5A-B). Assuming translation-invariance of connectivity strength such that it depends only on the distance between neurons’ preferred features, the network activity is symmetric under the translation of stimulus location.

**Figure 5.**
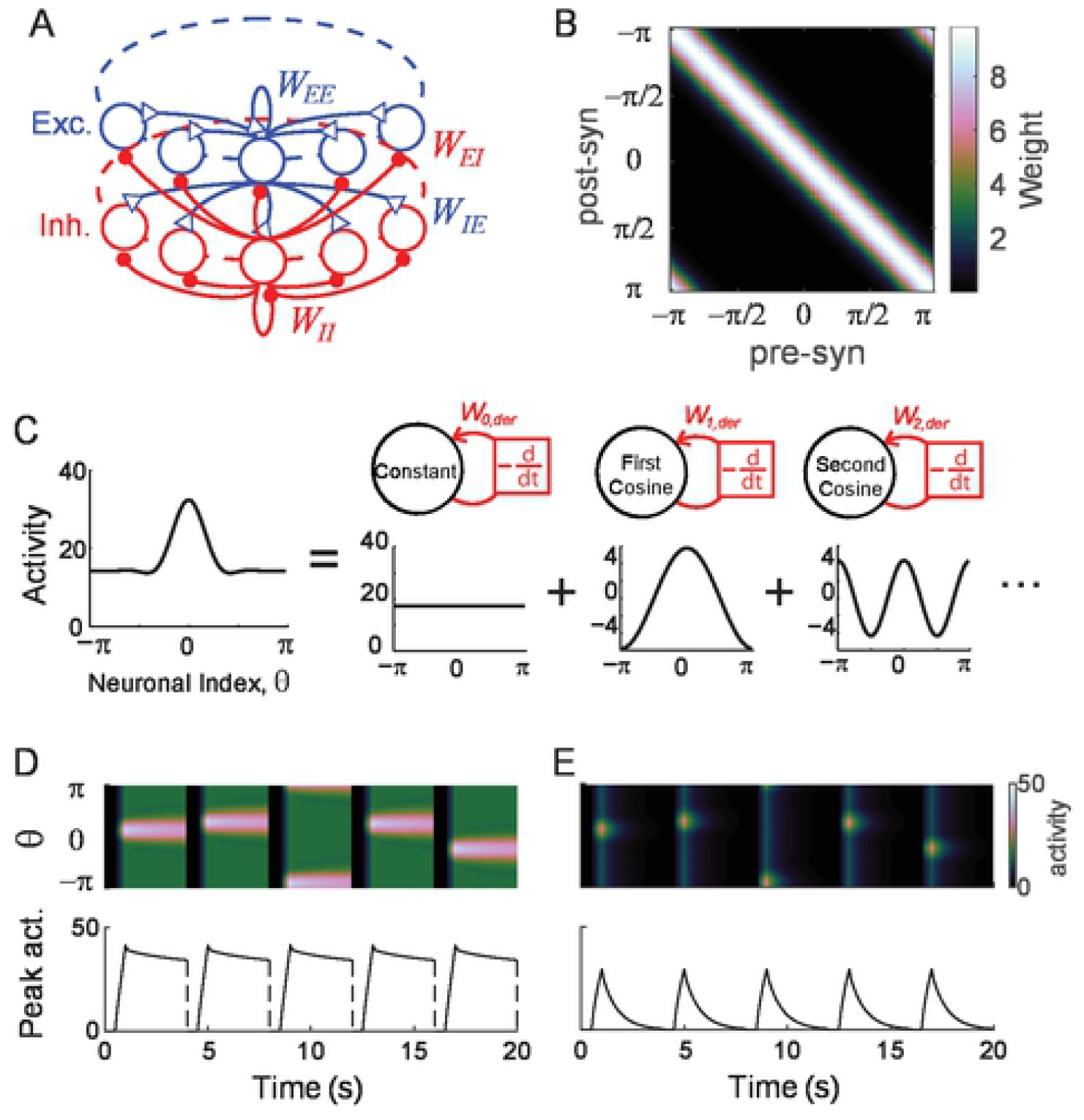
Location-coded persistent activity and its disruption under perturbation of tuning. A: Schematics of the spatial structure of network for location-coded memory. We considered that both excitatory and inhibitory neurons are organized in a columnar structure where each column consists of neurons with a similar preferred feature of the stimulus. Blue and red represent excitatory and inhibitory connections, respectively. B: Example connectivity matrix showing symmetry under translation. We considered the stimulus feature neurons encode the spatial information during the delay period, which lies on a circle, represented by θ ranging between -π and π. We assumed translation-invariance with the synaptic strengths depending only on the difference between preferred features of post and presynaptic neurons. C: Decomposition of spatially patterned activity into Fourier modes. Under translation-invariance, the activity can be decomposed into Fourier modes, and with strong negative derivative feedback, the dynamics of each mode become independent. D: Location-coded persistent activity under E-I balance. The activity during five consecutive trials was shown where the center of input is shifted randomly, showing maintenance of the spatial pattern of activity (upper panel) as well as elevated persistent activity at the stimulation center (lower panel). E: Disruption of persistent activity under 10% global perturbation in the E-to-E connection.

Under translation-invariant connectivity and activity patterns, dynamics can be analyzed through Fourier analysis ((12); Fig. 5C). For a large recurrent excitation and inhibition, the dynamics of Fourier modes are approximately independent of each other, each of which is analogous to the dynamics of one homogeneous population. Thus, the condition for negative derivative feedback in each Fourier mode is similar to the rate-coded network - slower recurrent excitation with the same condition on the synaptic time constants as in the homogeneous case, and balanced recurrent excitation and inhibition of that mode represented in terms of the Fourier coefficients of the synaptic strengths.

With similar balanced tuning conditions for the location-coded persistent memory, the perturbation to the synaptic connections leads to a similar disruption in the activity as in the rate-coded network (Fig. 5D-E). We first considered the multiplicative scaling down of all E-to-E connections, called a global perturbation (Fig. 5E). This leads to imbalanced excitation and inhibition and decay of activity in all Fourier modes. Note that the translation-invariant property is maintained under the global perturbation of the connectivity. Thus, the activity pattern is still symmetric for different stimulus locations despite its rapid decay to the baseline compared to the unperturbed case.

### Effects of plasticity under global perturbation

Next, we examine whether differential plasticity and homeostatic plasticity can recover the balance tuning condition for a spatially structured network. We assumed that the stimulus location across different trials is uniformly distributed and changes fast enough compared to the speed of the synaptic plasticity. Even if the connectivity is translation-invariant initially, stimulus at a particular location can lead to asymmetrical updates in the synaptic connections. Such asymmetrical updates can be mediated by slow learning and random stimulus locations having uniform distribution (30).

Under the maintenance of translation-invariance, the differential rule was shown to recover persistent activity having spatial patterns (Fig. 6). Unimodal activity peaked at the stimulus location can be maintained at any location after the differential plasticity rule recovers the balance of excitation and inhibition (Fig. 6A-B). We quantified the ability to maintain location-coded persistent memory using the decoding accuracy of spatial information at the end of the delay period (Methods). Initially, after global perturbation, the decoding error became around one, indicating loss of spatial selectivity, but over the course of learning with differential plasticity, it becomes close to zero (Fig. 6C). In line with this, the time constant of decay of different Fourier modes was shown to prolong (Fig. S1). In the eigenvector decomposition of the connectivity matrix, eigenvectors corresponding to the leading eigenvalues were found to be similar to Fourier modes, which is a signature of preservation of translation-invariance ((40); Fig. S1). The ratios of associated eigenvalues increase to one, albeit the different speeds, suggesting the recovery of balance tuning condition in each mode (Fig. 6D).

**Figure 6.**
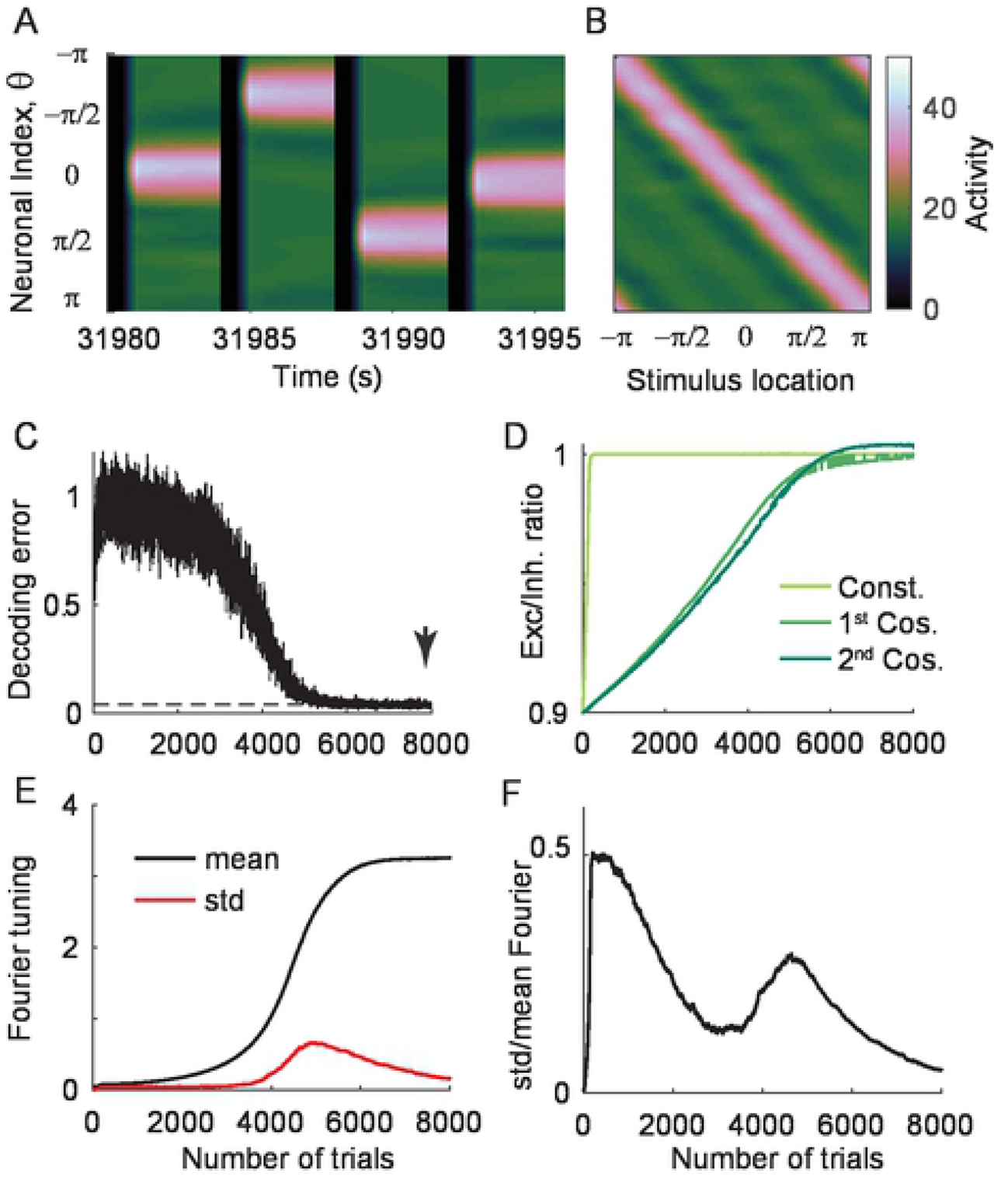
The effect of differential plasticity under weak global perturbation. A: Recovery of location-coded memory with differential plasticity under 10% global perturbation in the E-to-E connections. B: Activity pattern at the end of delay period after the recovery. With the connectivity frozen at trial 8000 (arrow in C), the spatial pattern of activity at the end of the delay period was shown for different stimulus locations. C: Improvement of decoding accuracy with learning. An individual trial refers to one memory task with a specific stimulus location. For each trial, we took the snapshot of activity as in B and quantified the decoding error using the population vector decoder (Methods). Dashed line indicates decoding error before perturbation D: Recovery of E-I balance for different Fourier modes. The eigenvector decomposition reveals the effective time constant of decay and recovery of E-I balance in different Fourier modes (Methods; Fig. S1). E: Mean (black) and standard deviation (red) of spatial selectivity across neurons quantified by the first Fourier component of each neuron’s tuning curve at the end of the delay period. F: Normalized standard deviation of Fourier tuning in (E) where its decrease with learning indicates recovery of translation invariance.

However, if translation-invariance breaks down, then Fourier analysis cannot be applied. This breakdown can occur either when the learning is too fast such that it cannot overcome asymmetry introduced by the different stimulus location at each trial or when the perturbation is too strong such that the activity of some neurons is stabilized to zero. Figure 7 shows the latter case – for stronger perturbation, the persistence of activity is recovered under differential plasticity, but the spatial pattern is fragmented by silent neurons (Fig. 7A-B). The decoding accuracy still improves during learning due to active neurons encoding spatial information (Fig. 7C and 7E).

**Figure 7.**
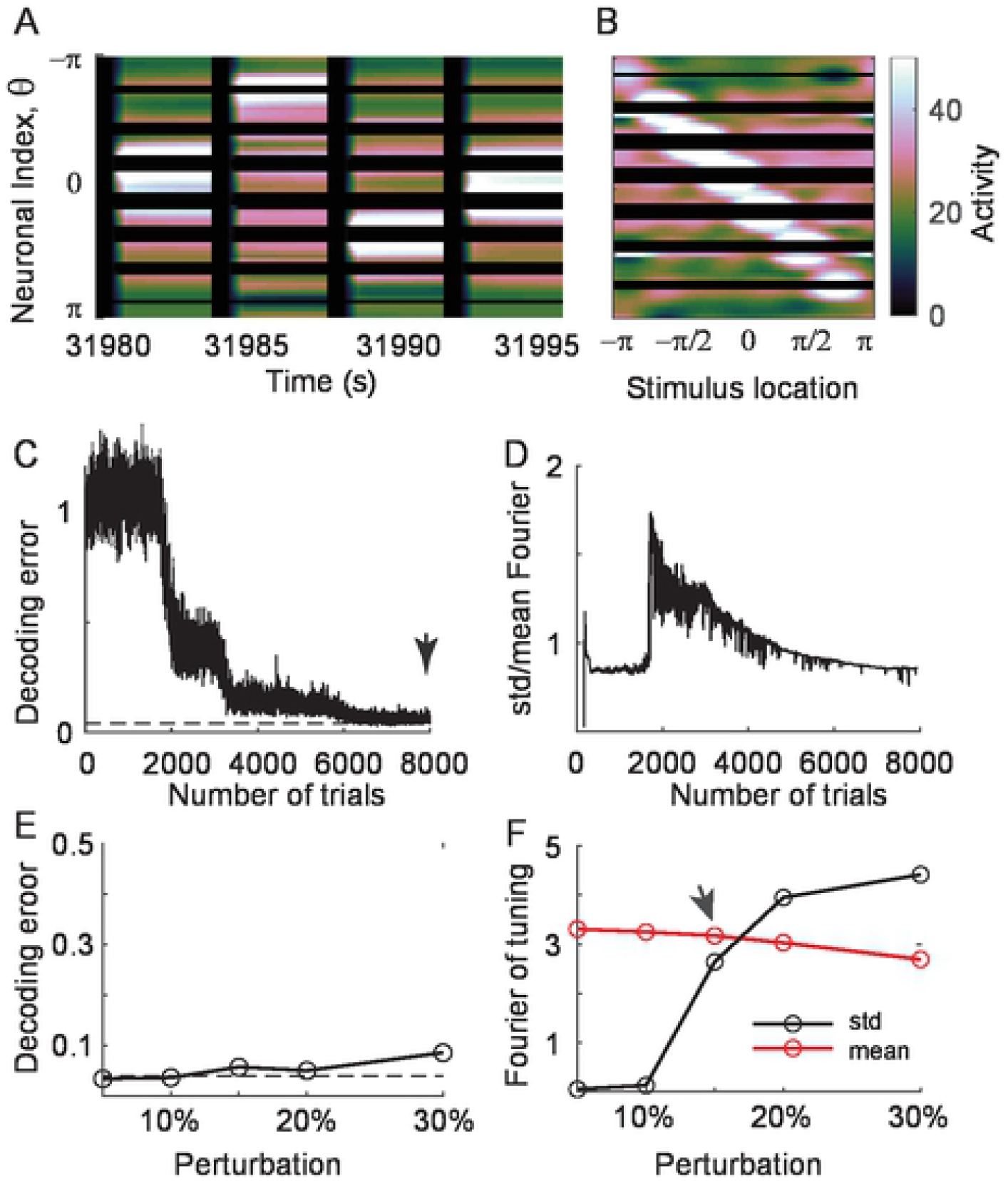
The effect of differential plasticity under larger global perturbation. A-B: Time course and tuning of activity after the recovery of E-I balance under differential plasticity (arrow in C), but 15% global perturbation was used. Differential plasticity stabilizes some neurons’ activity to zero, making them unresponsive to all stimulus locations. C: Improvement of decoding accuracy despite silent neurons due to compensation by neighboring active neurons. D: Normalized standard deviation of the first Fourier component over neurons, which remains high at the end of trials. E: Decoding errors under different strengths of perturbations. Dashed line indicates decoding error before perturbation. F: mean (black) and standard deviation (red) of spatial selectivity (F) under different strengths of perturbations.

For larger perturbation, the recovery of persistent activity and decoding accuracy is not uniform across different neurons as translation-invariance breaks down. To quantify this heterogeneity, we calculated the first Fourier component of the tuning curve of each neuron at the end of the delay period, representing its spatial selectivity, and obtained its mean and variance across neurons (Fig. 6E, 7D, and 7F; Method). Its mean increases with learning, indicating the increase of spatial selectivity with learning, shown for a broader range of perturbation (Fig. 6E and 7F). For a relatively weak global perturbation, the variance can transiently increase, reflecting an overall increase of activity level, but the normalized variance by the mean decreases for smaller perturbation with the translation-invariance maintained (Fig. 6E-F). However, for larger perturbation, such normalized variance is not reduced to zero even after decoding accuracy reaches its asymptote, indicating the breakdown of translation invariance (Fig. 7D and 7F).

While the maintenance of translation-invariance is not guaranteed under differential plasticity, homeostatic plasticity has been suggested to recover translation-invariance perturbed under cellular heterogeneity or other types of synaptic plasticity such as Hebbian learning (29). Indeed, the application of homeostatic learning rule to negative derivative feedback network recovers persistent unimodal activity at different locations (Fig. 8). As for differential plasticity, decoding accuracy improves with learning as the E-I balance is recovered (Fig. 8C and 8E). In contrast to differential plasticity, homeostatic plasticity achieved a low variance of spatial selectivity across neurons for a broader range of global perturbation, suggesting the maintenance of translation-invariance (Fig. 8D and F). As for a rate-coded memory network, homeostatic plasticity requires the input strengths to be tuned to match average delay activity to the target rate. However, the condition on the input strength can be mitigated with cellular or synaptic nonlinearity for location-coded persistent memory. The information can be decoded from the peak of the bump activity and is not sensitive to the amplitude of the bump.

**Figure 8.**
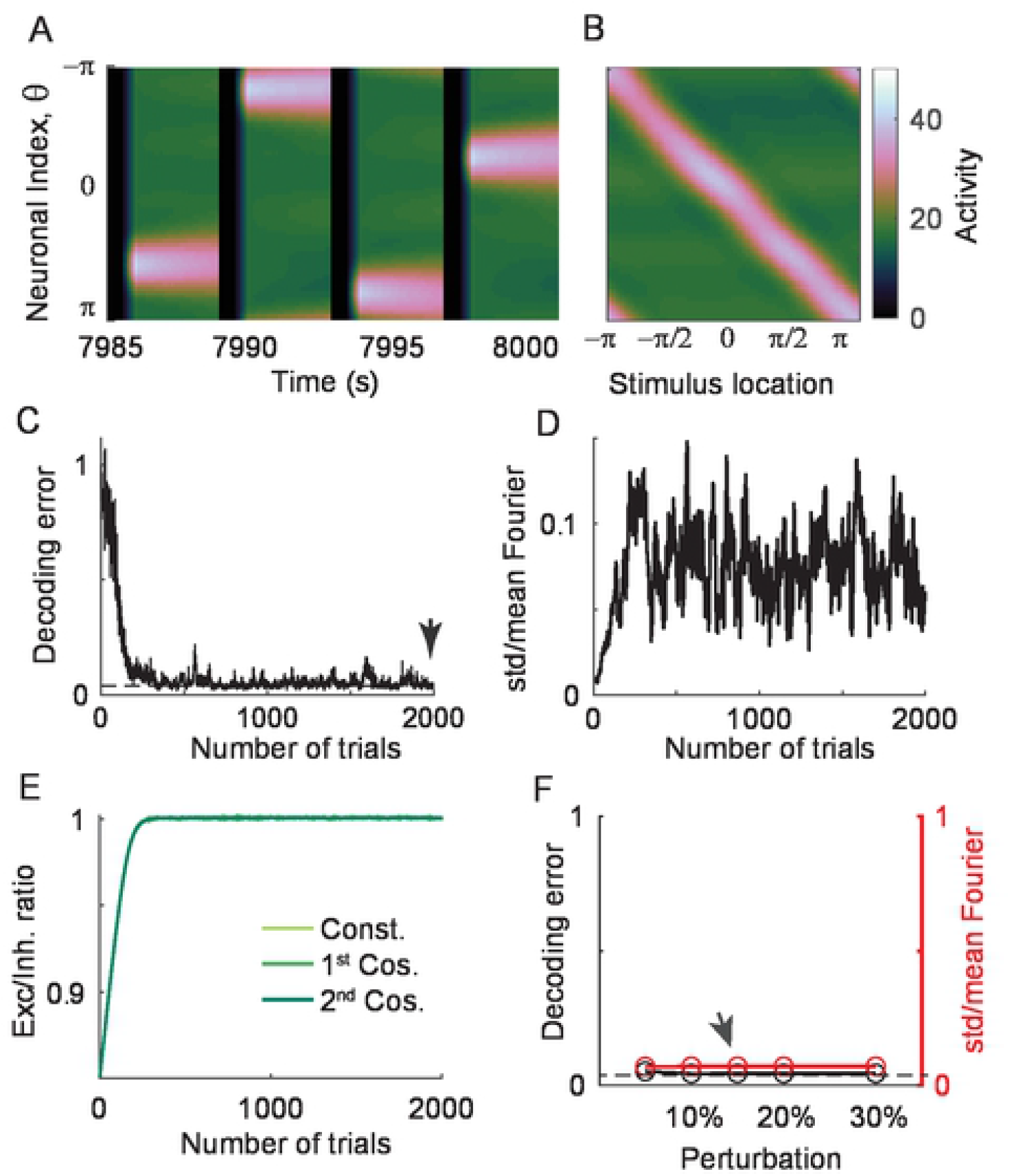
The effect of homeostatic plasticity under global perturbation. A-D: Same as Fig. 7A-D but with homeostatic plasticity, showing recovery of location-coded persistent activity and translation-invariance. E: Recovery of E-I balance in different Fourier modes shows the same recovery speed because homeostatic plasticity multiplicatively scales all afferent weights. F: Decoding errors (black) and normalized deviations of spatial selectivity (red) for different perturbation strengths. Dashed line indicates decoding error before perturbation.

### Effects of plasticity under local perturbation

We further investigated the effect of differential and homeostatic plasticity, where the balance of excitation and inhibition is locally perturbed. We considered two different types of local perturbations – first, postsynaptic perturbations, where synaptic strengths projected onto a particular group of neurons were perturbed (Fig. 9). For instance, this can be incurred by perturbation in NMDA receptors, which is considered to be prominent in the E-to-E connections (41). Mathematically, it is analogous to a row-wise perturbation in the E-E connectivity matrix (Fig. 9A). Another type of perturbation is the presynaptic one, where outgoing synaptic strengths are perturbed (Fig. 10). This perturbation can be caused by reducing transmitter release and is analogous to column-wise perturbation in the connectivity matrix (Fig. 10A). We considered a smooth bell-shaped perturbation assuming that the neurons with similar preferred spatial selectivity are clustered, and the effect of local perturbation dissipates across the clusters (42).

**Figure 9.**
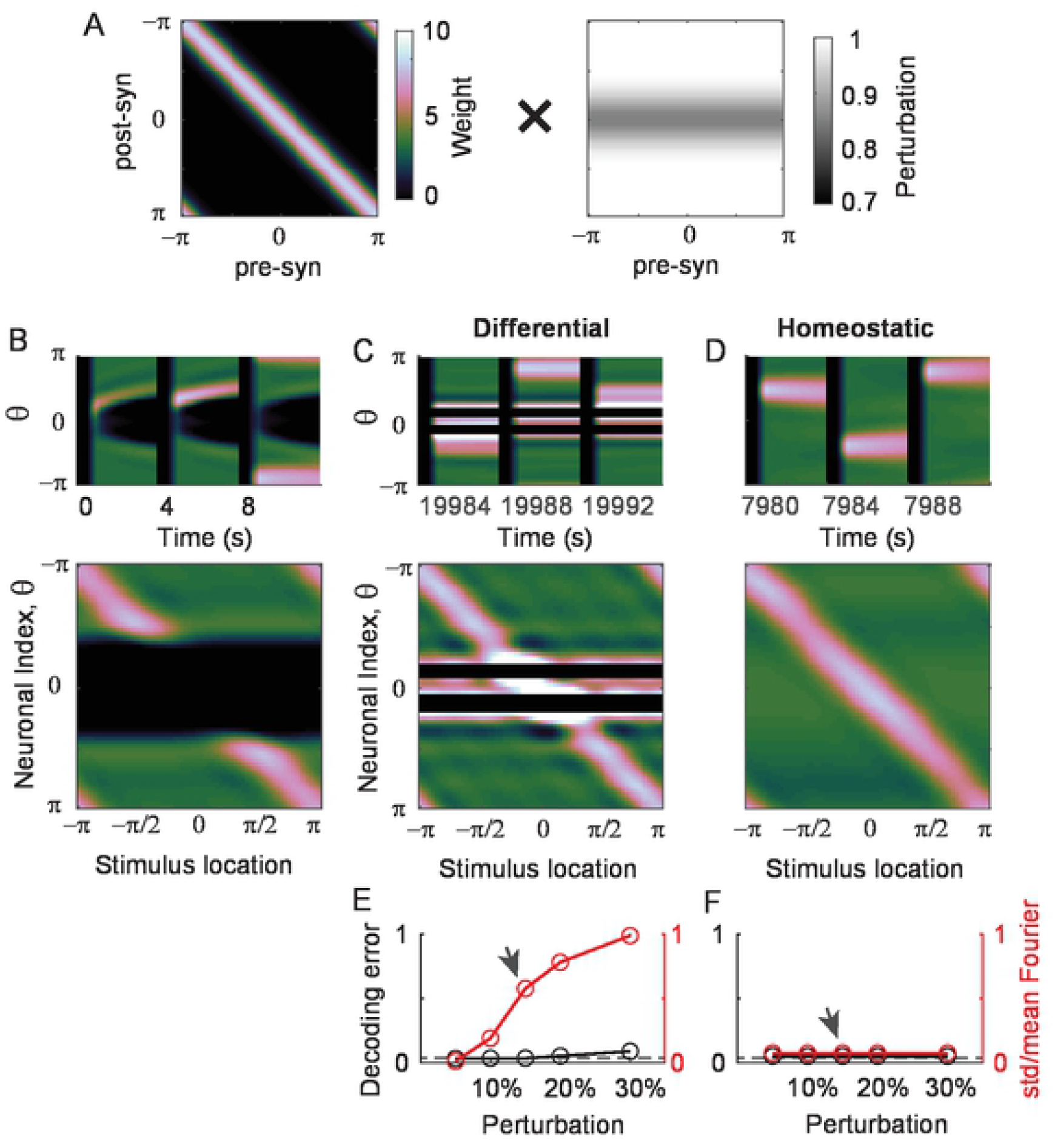
The effect of differential and homeostatic plasticity under postsynaptic perturbations. A: Schematics of postsynaptic perturbations where the rows of the connectivity matrix are multiplied by different scaling factors. Perturbation is centered at θ=0 and bell-shaped. B: Activity pattern under 15% post-synaptic perturbations before any plasticity. C-D: activity pattern shaped by the differential (C) and homeostatic (D) plasticity. E-F: Decoding errors (black) and normalized deviations of spatial selectivity (red) for different perturbation strengths after applying differential (E) and homeostatic (F) plasticity. Under differential plasticity, some neurons were silenced near the perturbation site (C), and the translation-invariance breaks down albeit with decent decoding performance (E). In contrast, homeostatic plasticity recovers the location-coded persistent activity for a broad range of postsynaptic perturbations (F).

**Figure 10.**
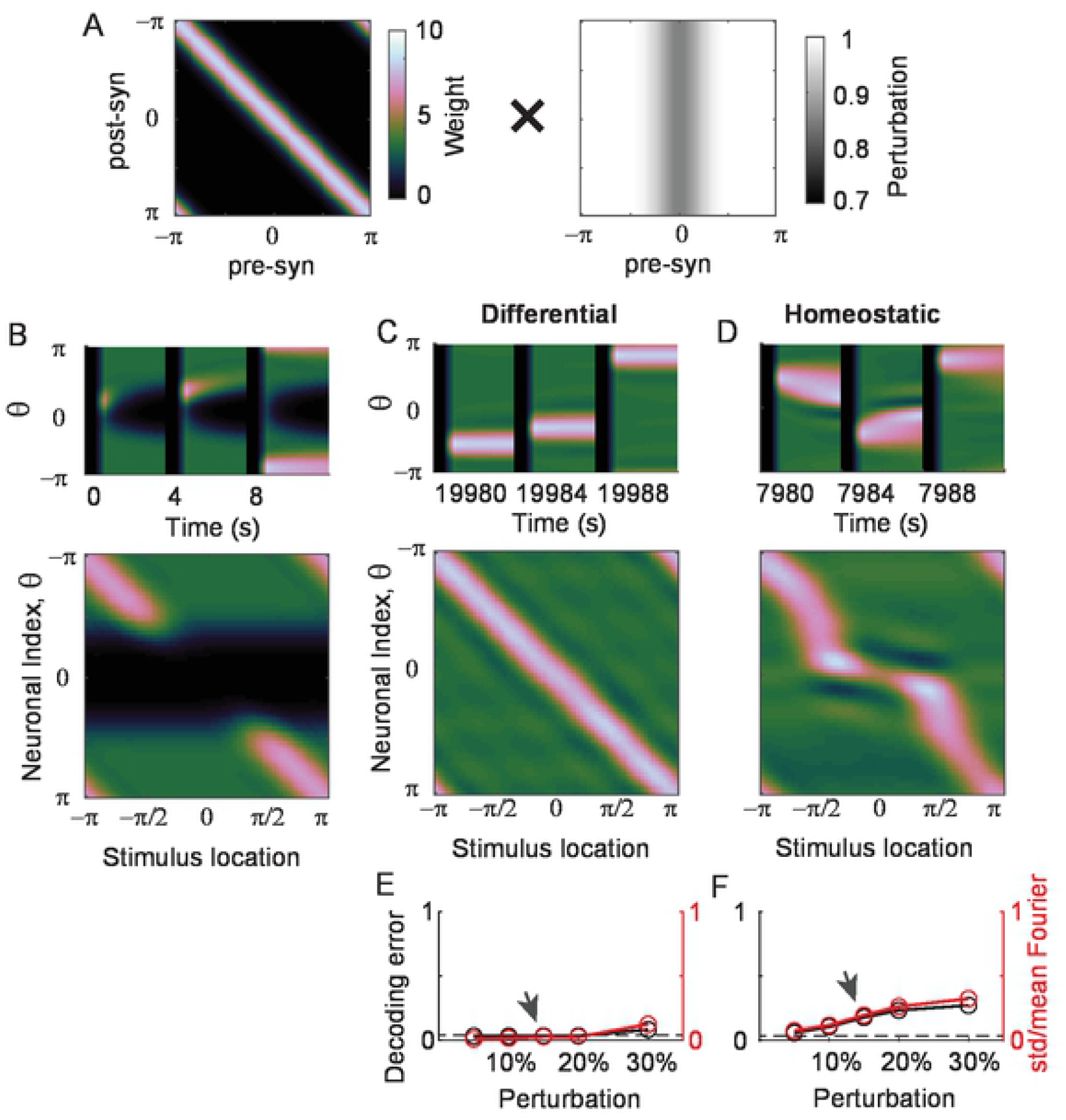
The effect of differential and homeostatic plasticity under pre-synaptic perturbations. A: Schematics of pre-synaptic perturbations where the columns of the connectivity matrix are multiplied by different scaling factors. B-F: Same as in Fig. 9B-F but under 15% pre-synaptic perturbation. Unlike postsynaptic perturbations, differential plasticity recovers persistent activity and translation-invariance (C, E). In contrast, with homeostatic plasticity, the activity pattern was distorted, resulting in worse decoding accuracy and translation-invariance for larger perturbations (D, F).

We first examined the effect of plasticity in postsynaptic perturbations. In negative derivative feedback models, the postsynaptic perturbation disrupts local E-I balance, leading to quick decay of activity in the vicinity of the perturbed site (Fig. 9B). Under relatively weak perturbation, both differential and homeostatic plasticity can recover E-I balance and the ability to maintain persistent activity at the perturbed site (Fig. 9E-F). However, when the perturbation becomes larger, differential and homeostatic plasticity show different recovery patterns as for the global perturbation (Fig. 9C-D). While homeostatic plasticity recovers both persistent activity and translation-invariance, differential plasticity persistently silences some neurons and cannot recover translation invariance. The fragmented spatial activity results in a high variance of spatial selectivity, breaking down translation-invariance, while the decoding accuracy is still good with compensation by higher activity at the vicinity of silent neurons (Fig. 9E). In contrast, homeostatic plasticity efficiently recovers translation-invariance caused by the overall reduction of synaptic strengths onto particular neurons as it multiplicatively scales those connections (Fig. 9F).

Next, we considered the effect of plasticity under presynaptic perturbations, which showed better performance of differential plasticity than homeostatic plasticity (Fig. 10). As in the postsynaptic perturbations, presynaptic perturbation causes activity at the perturbed site to decay because perturbation in outgoing synapses mostly affects the incoming synapses of neurons with similar spatial selectivity (Fig. 10B). Differential plasticity can recover persistent activity and translation-invariance for a broad range of presynaptic perturbation (Fig. 10C, 10E). On the other hand, homeostatic plasticity cannot stabilize persistent activity for relatively large perturbation, and the distortion of activity pattern is more substantial near the perturbed site (Fig. 10D). This is because presynaptic perturbation introduces an asymmetry in the synaptic strengths projecting onto neurons near the perturbed sites, which cannot be recovered through homeostatic plasticity that regulates the overall scaling of incoming synapses. Thus, although the average postsynaptic activity is recovered through increased excitability, the bump activity drifts towards instead of away from the perturbed site after learning, leading to a low decoding accuracy and breakdown of translation-invariance both (Fig. 10F).

### Effect of combining differential and homeostatic plasticity

As differential plasticity and homeostatic plasticity are effective in recovering persistent activity and translation-invariance under the different types of perturbations, we examined whether the combination of these two plasticities can utilize the advantage of both plasticities. Following the previous models considering the combination of Hebbian and homeostatic plasticity, we considered a multiplicative combination of two plasticities where differential plasticity replaces the Hebbian learning. The synaptic connection from neuron *j* to neuron *i* is expressed as a product of two variables, *W*_*ij*_ *= g*_*i*_*U*_*ij*_ with the dynamics of *g*_*i*_ and *U*_*ij*_ are given as

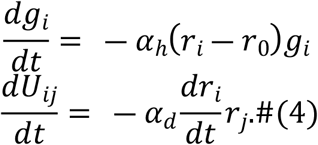

In the above equations, *g*_*i*_ reflects the homeostatic scaling, and *U*_*ij*_ evolves according to differential plasticity, with the learning rates given as α_h_ and α_d_, respectively.

We first examined the effect of combined plasticity under global perturbations. Although differential plasticity alone can lead to the silence of activity, homeostatic plasticity prevents it by boosting lower-than-target activity. Thus, combined plasticity could recover location-coded persistent activity and translation invariance for a broad range of perturbations (Fig. 11A and 11D). Note that such a recovery is sensitive to the learning rates of the plasticity such that the overall speed of both differential and homeostatic plasticity needs to be slow, but homeostatic one needs to be relatively fast. This is because too fast homeostatic learning leads to oscillation, yet it needs to be fast enough to prevent the breakdown of translation invariance introduced by differential plasticity.

**Figure 11.**
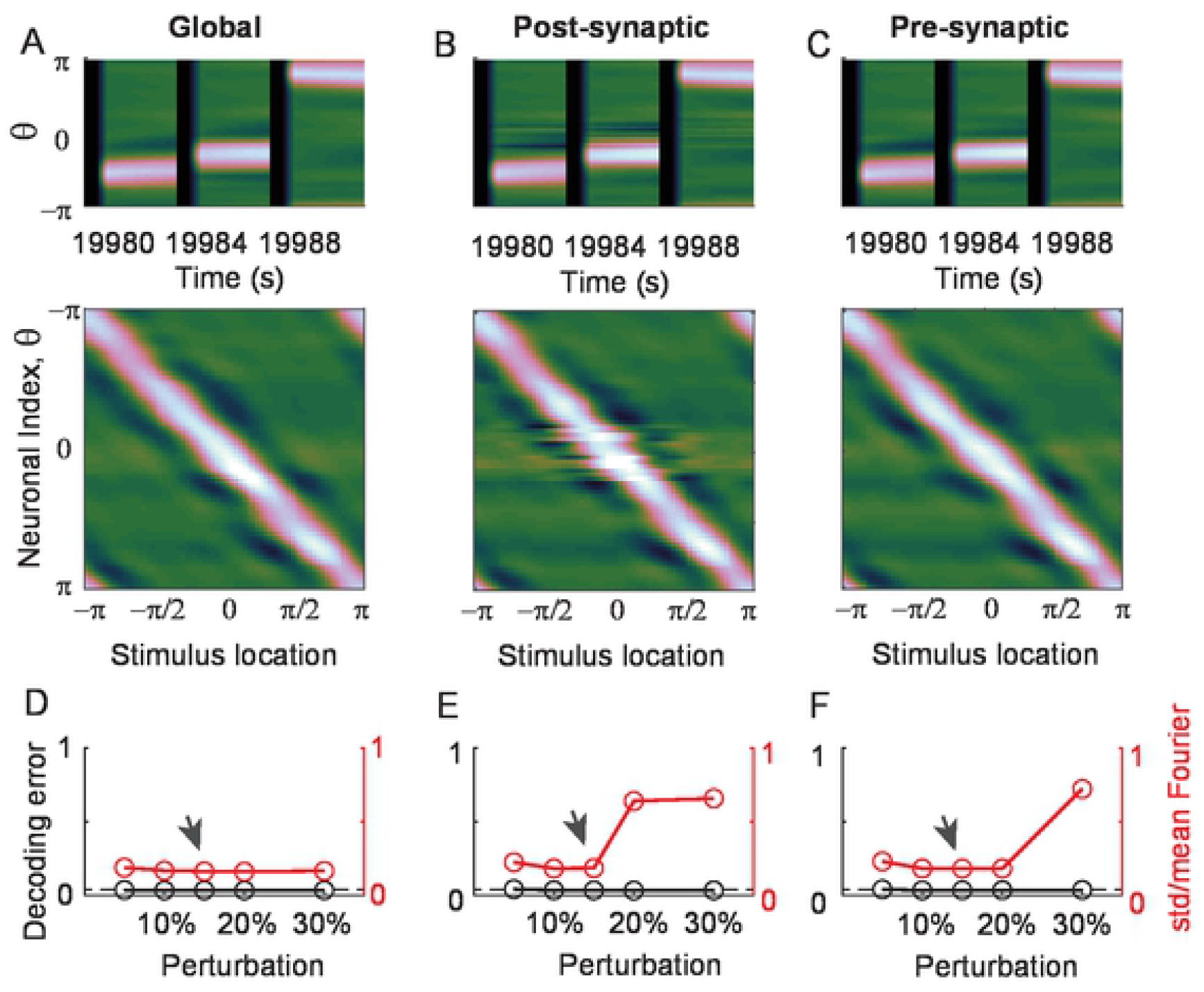
The effect of the combination of differential and homeostatic plasticity. A-C: Recovery of location-coded persistent activity under combined plasticity after 15% global (A), postsynaptic (B), and pre-synaptic perturbation (C). Activity pattern after 30% local perturbations is shown in Fig. S2. D-F: Decoding errors (black) and normalized deviation of spatial selectivity (red) for different perturbation strengths.

The combined plasticity also shows the compensation under both types of local perturbations (Fig. 11B-C, E-F). For a postsynaptic perturbation under which differential plasticity could not recover the translation-invariance, the combined one shows the extension of the recovery (Fig. 9C vs. 11B, 9E vs. 11E). The superiority of the combined one is similar for a presynaptic perturbation under which homeostatic plasticity could not recover both persistent activity and translation-invariance (Fig. 10D vs. 11C, 10F vs. 11F). Note that still, for a larger perturbation, the activity pattern can be distorted, and translation-invariance is not perfectly recovered (Fig. 11E-F, S2). However, the decoding accuracy is decent for a broad range for the combined plasticity.

## Discussion

In this work, we investigated the effects of local and unsupervised learning on the stabilization of persistent activity in two representative working memory models encoding analog values, namely, rate-coded and location-coded persistent memory. We examined the effects of differential plasticity and homeostatic plasticity by systematically varying the magnitude and form of perturbations in synaptic connections. Consistent with the findings of previous works, differential plasticity alone was enough to recover a graded-level persistent activity in a homogeneous population (11,18). On the other hand, recovery by homeostatic plasticity requires the tuning of learning parameters. For the maintenance of spatially structured persistent activity, differential plasticity could recover persistent activity, but its pattern can be irregular for different stimulus locations. On the other hand, homeostatic plasticity shows robust recovery of translation-invariance against particular types of synaptic perturbations, such as perturbations in incoming synapses onto the entire or local populations, which are similar to the inhomogeneity of neuronal gain considered previously (29). However, homeostatic plasticity was not effective against perturbations in outgoing synapses from local populations. Instead, combining it with differential plasticity recovers the location-coded persistent activity for a broader range of perturbations.

Persistent activity sustained in the absence of external stimuli has been suggested as a signature of static attractor dynamics. While most attractor models are based on positive feedback mechanisms, we considered negative derivative feedback mechanisms with two advantages to investigate the effect of synaptic plasticity. First, a negative derivative feedback network has similar tuning conditions in both the rate-coded and location-coded persistent memory with the same condition on the balanced excitation and inhibition and additional symmetry of translation-invariance for location-coded memory (12,33). Thus, the analysis of the effect of synaptic plasticity in a relatively simple rate-coded network could be extended to that in a location-coded network. Second, the negative derivative feedback model is less dependent on a specific form of intrinsic nonlinearity of neurons, so graded perturbation causes a graded change in the network’s behavior. On the other hand, intrinsic nonlinearity plays a critical role in positive feedback networks, which leads to additional complexity in systematically investigating the effect of synaptic plasticity (43) (but see (44,45)). However, note that our main findings may still be valid regardless of the specific underlying mechanism of working memory. For rate-coded persistent activity, previous works have shown that differential plasticity recovers the tuning condition of the positive feedback mechanism (17,18). We tested two plasticity rules on the location-coded memory based on positive feedback and verified that homeostatic plasticity is effective against post-but not presynaptic perturbation; however, differential plasticity can effectively stop drift but may lead to an irregular pattern, and combination of both provides a partial remedy (Fig. S3).

Stable memory formation under the mixture of different forms of synaptic plasticity has been proposed previously, mainly for discrete attractor networks (35,36,46). In these studies, Hebbian synaptic plasticity has been suggested to form auto-associative memory guided by external inputs. To prevent instability caused by Hebbian learning, compensatory mechanisms, such as homeostasis or short-term plasticity, were required, which must act on a timescale similar to that of Hebbian learning (47). Our work also suggests synergistic interplay between different types of plasticity, differential, and homeostatic plasticity, in particular for stabilizing location-coded persistent memory. However, we note that differential plasticity alone is stable. The role of homeostatic plasticity is to support translation-invariance in a ring-like architecture of recurrent connections (29,30). Thus, the fast dynamics of homeostatic plasticity are not required, and excessively fast dynamics can be detrimental due to oscillatory instability. The interplay between anti-Hebbian learning and activity-dependent synaptic scaling has been proposed for rate-coded persistent memory (48), where the anti-Hebbian rule itself stabilizes the network activity and no fast homeostasis is required, as in our work.

In this work, we assumed the existence of synaptic plasticity only during the delay period. However, differential plasticity might make the network “unlearn” if it operates the same way during the stimulus period as in the delay period because the activity rise during that time would be interpreted as positive drift by the plasticity. It is thus essential to constrain derivative-driven learning only during a period in which the activity ought to be stabilized. In oculomotor literature, such as (17), this is done by filtering fast-changing activity that is potentially related to burst of the saccadic signal. When we consider the biological implementation of differential plasticity, there is an alternative way that this can happen. Xie and Seung (2000), motivated by (23), showed that spike-timing-dependent plasticity (STDP) is intrinsically sensitive to the time derivative of activities, and it can be approximated by the differential plasticity considered in our work when the overall potentiation and depression are balanced (18). Alternatively, Nygren et al. (2019) showed that similar differential plasticity could be implemented through cancelation with a delayed feedback signal analogous to the derivative feedback (19). In both cases, the derivative is approximated for slowly changing neural activity, and higher frequency changes are filtered out. On the other hand, continuous learning with homeostatic plasticity may require the adjustment of learning parameters because the long-term average firing rates of neurons must reflect activity during the entire session.

Constraining activity drifts of individual neurons might require stricter conditions than what is required to achieve stable coding of information during the memory period. While traditional experimental work identified memory neurons that showed elevated persistence firing with stimulus selectivity (42), recent population-level analysis revealed the stable readout of information across various time points despite the diverse temporal dynamics of individual neurons (49,50). Such dynamic activity in individual neurons may reflect activity in the downstream population that combines stimulus-encoding persistent activity and time-varying activity, possibly reflecting time information (51,52). On the other hand, memory networks themselves can allow time-varying activity such that attractor dynamics are formed along with the particular activity pattern or mode, while allowing temporal fluctuation along with other modes (50,53). For the latter, synaptic plasticity based on the global error signal has been suggested, which can be a self-supervised signal, such as a drift in the readout activity (20) or a difference from the target signal (54). Note that the resulting form of synaptic plasticity is similar to differential plasticity, where the activity drift of individual neurons in differential plasticity is replaced with the global error signal. Homeostatic processes, such as intrinsic plasticity, inhibitory plasticity, and synaptic scaling, have also been proposed to elongate memory traces in the presence of dynamic activity (48,55). In these works, the memory is maintained by a network with minimally structured connectivity, and the sensitivity to learning parameters has not been analyzed.

Overall, our work demonstrates how unsupervised learning can mediate fine-tuning conditions for working memory implementing continuous attractors. It aligns with previous works emphasizing the role of unsupervised learning to generate a basis of activity patterns and dynamics underlying cognitive functions (56–58). While we focused on unsupervised learning rules regularizing temporal patterns in the absence of input, they can be combined with other learning rules that can act under the guidance of external inputs and may make memory networks robust for a broader range of perturbations. Also, we considered perturbation and synaptic plasticity only in a specific connection, recurrent E-to-E connections, but the plasticity of other connections, such as inhibitory plasticity (59–61), has been suggested to tune network homeostasis and EI balance. Given the importance of balance and homeostasis in memory circuits, further investigation is needed to examine the effect of unsupervised plasticity on various synapses. Also, to understand how the learning parameters of these plasticity rules match with neural activity, a detailed investigation of the underlying biophysical mechanisms needs to be done, possibly in models involving multiple subcellular compartments.

## Methods

All codes are available at https://github.com/jtg374/NDF_ringNet_plasticity

### Simple rate model for a homogeneous population

In this section, we show the derivation of a one-dimensional differential equation in Eq. 1 (see more biological structure and conditions in Lim & Goldman, 2013). For this, we considered one homogeneous population receiving recurrent excitation and inhibition with different kinetics, described by three-dimensional differential equations

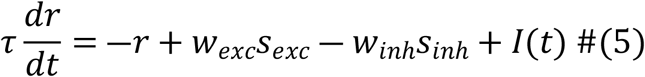

where three dynamic variables are firing rate *r*, recurrent excitatory currents *s*_*exc*,_ and recurrent inhibitory currents *s*_*inh*_. We assumed that *s*_*exc*_ and *s*_*inh*_ are low-pass filtered *r* with time constants *τ*_*exc*_ and *τ*_*inh*_, respectively, and *s*_*exc*_ ≈ *r* when *r* hardly changes.

With *s*_*exc*_ – *s*_*inh*_ ≈ -(*τ*_*exc*_ - *τ*_*inh*_)*dr/dt*, the above equation can be approximated as a one-dimensional differential equation, given as

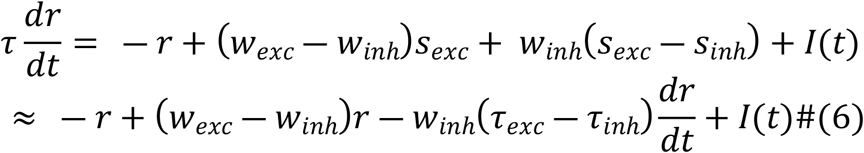

With *w*_*exc*_ *-w*_*inh*_ and *w*_*inh*_(*τ*_*exc*_ - *τ*_*inh*_) denoted by *w*_*net*_ and *w*_*der*_, Eq. 6 is the same as Eq. 1. Such one-dimensional approximation allows analytic investigation on the effects of differential plasticity and homeostatic plasticity in Eq. 2 and Eq. 3.

In Eq. 5, when we normalized *w*_*exc*_ with *w*_*inh*_ denoted as *w = w*_*exc*_*/w*_*inh*_ and assumed *w*_*inh*_ is large such that *w*_*der*_ >> *τ*, Eq. 6 becomes

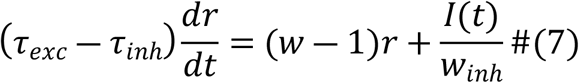

During the delay period with no external input *I(t)*, the dynamics with the differential plasticity in Eq. 2 becomes

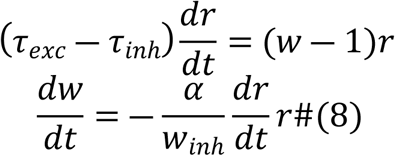

Thus, varying *w*_*inh*_ has the same effect as changing the learning speed α as illustrated in Fig. 2B,C. On the other hand, the dynamics with homeostatic plasticity in Eq. 3 becomes

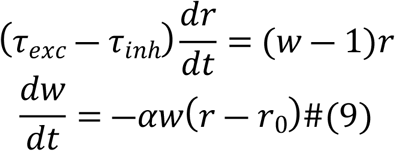

That is, changing *w*_*inh*_ has no effect on the recovery speed for homeostatic plasticity.

In Equations 1-3, we set *τ* and *τ*_*exc*_ *-τ*_*inh*_ to be unit time constant 1, and initial *w*_*exc*_, *w*_*inh*_ *and w*_*der*_ are set to be 500. *I(t)* is a step function, giving the input for 50 unit time with its strength randomly distributed as 0 and 1000 during the learning, and three representative traces of *r(t)* for *I(t)* = 250, 500 and 1000 were shown before and after learning. For the differential plasticity, the learning speed α is 0.01. For homeostatic plasticity, α is 2×10^−5^ in Fig. 3 and Fig. 4a and b and 0.001 in Fig. 4c. r_0_ is 50 in Fig. 3 and Fig. 4c, and 80 and 25 in Fig. 4a, and b.

### Spatial structured network model for location-coded persistent activity

Following Lim & Goldman 2014, we considered a network organized in a columnar architecture for spatial working memory with the equations describing the dynamics given as

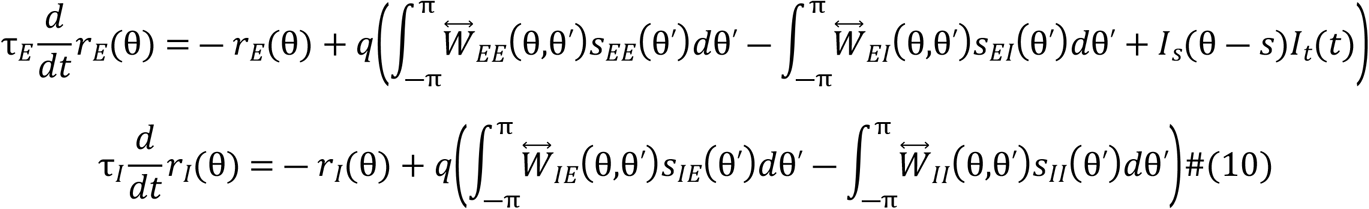

where subscripts *E* and *I* represent excitatory and inhibitory populations, respectively. The activity and the connectivity were indexed by their preferred spatial feature, θ, ranging between [-π,π). τ_E_ and τ_I_ are the time constants and *q(·)* is the input-output transfer function, which is the rectified linear function given as *q(x) = x* for *x > 0* and otherwise, 0. ^↔^W_ij_ (*i, j* = E or I) is the synaptic weight and before perturbation, it was taken to be translation-invariant and Gaussian-shaped as

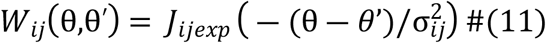

As in the homogeneous case, 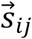 (*i, j* = E or I) represents the synaptic variables whose dynamics is given as

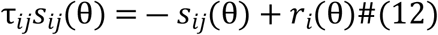

Importantly, the excitatory-to-excitatory (E-to-E) time constant needs to be much larger than other synapses’ to make derivative feedback happen (Lim & Goldman 2013). Detailed parameters used in the simulation will be given in the section “Table of parameters”.

I_s_(θ-s) and I_t_(t) represent the spatial and temporal profiles of external stimulus where s is the center of the stimulus location. I_s_(θ) also has a Gaussian shape as

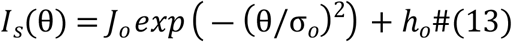

I_t_(t) is a pulse function smoothed by a low-pass filter with time constant τ_o_

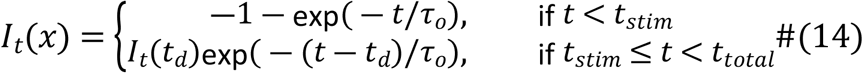

### Perturbation and plasticity model

We considered three types of perturbations in the E-to-E connections. For the global perturbation, ^↔^W_EE_ was set to be

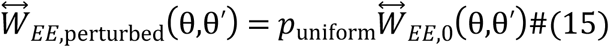

Postsynaptic perturbation corresponds to a row-wise change as

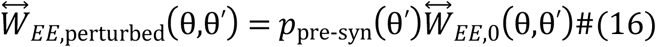

Similarly, presynaptic perturbation corresponds to a column-wise change as

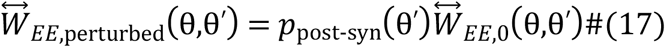

Where p(θ) is a smooth function of θ, given as a Gaussian function

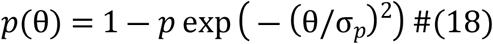

To recover the persistent activity, we considered two types of plasticity, differential and homeostatic plasticity, described as

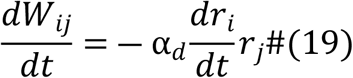

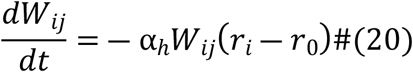

where α_d_ and α_h_ represents the learning rate of differential and homeostatic plasticity. In the combined one in Eq. 4, ^↔^W_ij_ in Eq. 2 and Eq. 3 are replaced by *U*_*ij*_ and *g*_*i*_, respectively. The plasticity is only applied in the delay period, and to minimize the effect of the residual stimulus, we also gated the plasticity with a factor 1-I_t_(t), though it does not make much difference if we don’t add it.

### Quantify E-I balance through eigenvalue decomposition

In Figure 6D and 8E, we quantify the recovery of EI balance by taking the eigenvalues of the weight matrices. When translation-invariance is preserved, the values of both E-to-E matrix and other weight matrices will approximately be the Fourier components of the matrices and the tuning conditions for n-th Fourier modes become

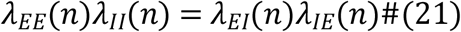

where *λ*_*ij*_(*n*) is the *n*-th eigenvalue of ^↔^W_ij_. In Fig. 6D and 8E, we did the eigenvector decomposition of the weight matrix ^↔^W_EE_ and found the eigenvectors resembles Fourier modes and calculate E-I balance ratio in each mode from the corresponding eigenvalues.

### Decoding error

We quantified the network’s memory performance by decoding the stimulus at the end of the delay. Because we used a deterministic simulation, we modeled the noise post-hoc with Poisson random number generator, assuming the spike generation is random and independent across neurons. For each neuron, we multiplied its firing rate (in Hz) by 0.2 and used the product as the mean of the Poisson random number to model its spike count in 200ms. We then decoded the location from the simulated spike count *n*_*θ*_ with a simple population-vector decoder:

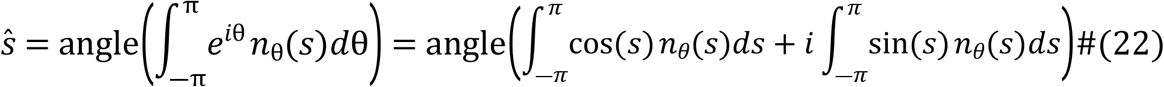

The error is quantified by the cosine distance between the decoded location and true stimulus:

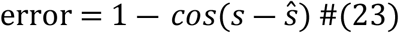

For each stimulus, the random generation of spike counts was repeated 20 times and averaged. We quantified the average error across all stimulus locations by freezing the network connectivity at each trial. After perturbation, when there is no spatial selectivity at the end of the delay, ŝ would be uniformly distributed, and the average error would be one, while if the spatially patterned activity is persistent with no drift, the decoding error would be close to zero. In Figure S3, we divided the stimulus locations into eight groups and visualized the average error within groups to emphasize that local perturbation affects decoding accuracy differently depending on stimulus locations.

### Spatial selectivity and translation invariance

The spatial selectivity of each neuron was quantified by calculating the first Fourier component of its tuning curve given as

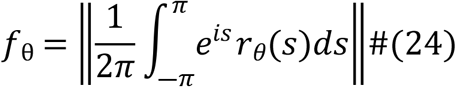

where r_θ_(s) is the activity of the neuron that is selective to θ at the end of the delay period of a trial stimulated at s.

We calculated the mean and standard deviation across neurons.

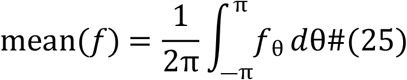

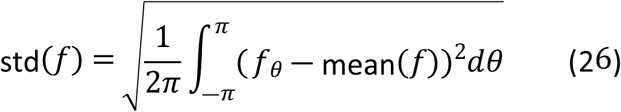

The normalized std (std/mean) was used to quantify translation-invariance in Fig. 6-11.

### Table of parameters for spatially structured network

| Parameter | Description | Value |
| --- | --- | --- |
| $N$ | Number of population in each group | 64 |
| $\tau_E$ | Time constant of excitatory neurons | 20 |
| $\tau_I$ | Time constant of inhibitory neurons | 10 |
| $\tau_{EE}$ | Time constant of E-to-E synapses | 100 |
| $\tau_{EI}$ | Time constant of I-to-E synapses | 10 |
| $\tau_{IE}$ | Time constant of E-to-I synapses | 25 |
| $\tau_{II}$ | Time constant of I-to-I synapses | 10 |
| $\tau_o$ | Time constant of external stimulus | 100 |
| $J_{EE}$ | Amplitude of E-to-E synaptic weight | 100 |
| $J_{EI}$ | Amplitude of I-to-E synaptic weight | 100 |
| $J_{IE}$ | Amplitude of E-to-I synaptic weight | 200 |
| $J_{II}$ | Amplitude of I-to-I synaptic weight | 200 |
| $J_o$ | Amplitude of external stimulus | 270 |
| $\sigma_{EE}, \sigma_{IE}$ | Width of excitatory synaptic connections | $0.2\pi$ |
| $\sigma_{EI}, \sigma_{II}$ | Width of inhibitory synaptic connections | $0.1\pi$ |
| $\sigma_o$ | Width of stimulus | $0.25\pi$ |
| $h_0$ | Baseline of stimulus | 200 |
| $p$ | 1 - perturbation strength | vary |
| $\alpha_d$ | Learning rate of differential rule | 1e-5 |
| $\alpha_h$ | Learning rate of homeostatic rule | 2e-8 |
| $r_0$ | Target firing rate of homeostatic rule | 20 |
| $t_{stim}$ | Stimulation duration | 500 |
| $t_{total}$ | Stimulation plus delay period | 3500 |

## Acknowledgments

The authors acknowledge the support from NYU-ECNU Institute of Brain and Cognitive Science

## Supplementary Information

**Figure S1, related to Fig. 6, Elongation of time constant associated with each eigenvector similar to Fourier modes under differential plasticity**.

A: Time scale of each Fourier mode. For each Fourier mode, a time constant was estimated by projecting population activity onto a sinusoid of different frequencies and fitting the time course with exponential decay. The negative reciprocals of these time constants have good correspondence with the eigenvalues shown in Fig 6D. B: Eigenvectors related to eigenvalues in Fig. 6D during the evolution of learning dynamics. The real part of the eigenvectors corresponding to the first, third, and fifth leading eigenvalues (even ones omitted because of redundancy) is plotted. The shape of the eigenvectors is close to sinusoids, suggesting preservation of translation-invariance.

**Figure S2, related to Fig. 11. The effect of combined plasticity under larger local perturbation**.

A-B: same as Fig 11 B-C, but under 30% local perturbation, as indicated by arrowhead in C-D. C-D: copy of Fig 11E-F. For larger local perturbation, translation-invariance breaks down while decoding errors are low. The distortion of activity patterns is dissimilar to that only with differential plasticity because homeostatic plasticity keeps neurons from falling silent in some trials but not in others.

**Figure S3. The effect of differential, homeostatic, and combined plasticity rules in positive feedback network under 30% pre-synaptic perturbation**.

A-C: Decoding errors for eight stimulus groups during the recovery. Under the local perturbation, translation-invariance breaks down in the positive feedback models, and the bump activity tends to drift as in a negative derivative feedback network. The effects under pre-synaptic perturbations were only shown for simplicity. D: Activity pattern with homeostatic plasticity at around trial 4000 (arrows in A). E, F: Activity pattern with differential and combined plasticity at around trial 2000 (arrows in B, C). As in the negative derivative feedback network, homeostatic synaptic plasticity is not effective under pre-synaptic perturbation (D). In contrast, differential plasticity effectively stops the drift (E), as well as the combined plasticity (F). Note that the activity patterns under the differential (B) and combined plasticity (C) start to break down around 3000 trials after the recovery. This breakdown may arise from implementing a raw dr/dt in differential plasticity, which is sensitive to a slight change of activity pattern as it evolves from stimulus-evoked patterns to the stereotypical delay activity patterns.

## References

1. Knierim JJ, Zhang K. Attractor dynamics of spatially correlated neural activity in the limbic system. Annu Rev Neurosci. 2012;35:267–85.

2. Goldman MS, Compte A, Wang XJ. Neural Integrator Models. In: Encyclopedia of Neuroscience. Elsevier Ltd; 2009. p. 165–78.

3. Durstewitz D, Seamans JK, Sejnowski TJ. Neurocomputational Models of Working Memory. Nat Neurosci. 2000;3(11s):1184–91.

4. Aksay E, Baker R, Seung HS, Tank DW. Correlated Discharge among Cell Pairs within the Oculomotor Horizontal Velocity-to-Position Integrator. J Neurosci. 2003;

5. Yoon K, Buice MA, Barry C, Hayman R, Burgess N, Fiete IR. Specific evidence of low-dimensional continuous attractor dynamics in grid cells. Nat Neurosci. 2013 Aug 14;16(8):1077–84.

6. Wimmer K, Nykamp DQ, Constantinidis C, Compte A. Bump attractor dynamics in prefrontal cortex explains behavioral precision in spatial working memory. Nat Neurosci. 2014;17(3):431–9.

7. Brody CD, Romo R, Kepecs A. Basic mechanisms for graded persistent activity: discrete attractors, continuous attractors, and dynamic representations. Curr Opin Neurobiol. 2003 Apr;13(2):204– 11.

8. Seung HS. How the brain keeps the eyes still. Proc Natl Acad Sci U S A. 1996;93(23):13339–44.

9. Koulakov AA, Raghavachari S, Kepecs A, Lisman JE. Model for a robust neural integrator. Nat Neurosci. 2002;5(8):775–82.

10. Goldman MS, Levine JH, Major G, Tank DW, Seung HS. Robust Persistent Neural Activity in a Model Integrator with Multiple Hysteretic Dendrites per Neuron. Cereb Cortex. 2003;

11. Lim S, Goldman MS. Balanced cortical microcircuitry for maintaining information in working memory. Nat Neurosci. 2013 Sep 18;16(9):1306–14.

12. Lim S, Goldman MS. Balanced Cortical Microcircuitry for Spatial Working Memory Based on Corrective Feedback Control. J Neurosci. 2014;34(20):6790–806.

13. Itskov V, Hansel D, Tsodyks M. Short-term facilitation may stabilize parametric working memory trace. Front Comput Neurosci. 2011 Oct 24;5.

14. Seeholzer A, Deger M, Gerstner W. Stability of working memory in continuous attractor networks under the control of shortterm plasticity. Burak Y, editor. PLoS Comput Biol. 2019 Apr 19;15(4):e1006928.

15. Arnold DB, Robinson DA. A neural network model of the vestibulo-ocular reflex using a local synaptic learning rule. Philos Trans R Soc Lond B Biol Sci. 1992;337(1281):327–30.

16. Major G, Baker R, Aksay E, Mensh B, Seung HS, Tank DW. Plasticity and tuning by visual feedback of the stability of a neural integrator. Proc Natl Acad Sci U S A. 2004;101(20):7739–44.

17. MacNeil D, Eliasmith C. Fine-tuning and the stability of recurrent neural networks. Vasilaki E, editor. PLoS One. 2011 Sep 27;6(9):e22885.

18. Xie X, Seung HS. Spike-based learning rules and stabilization of persistent neural activity. In: Solla SA, Leen TK, Müller K, editors. Advances in Neural Information Processing Systems. MIT Press; 2000. p. 199–205.

19. Nygren E, Ramirez A, McMahan B, Aksay E, Senn W. Learning temporal integration from internal feedback. bioRxiv. 2019;

20. Federer C, Zylberberg J. A self-organizing short-term dynamical memory network. Neural Networks. 2018 Oct 1;106:30–41.

21. Kosko B. Differential Hebbian learning. In: AIP Conference Proceedings. AIP; 1986. p. 277–82.

22. Der R, Martius G. Novel plasticity rule can explain the development of sensorimotor intelligence. Proc Natl Acad Sci. 2015 Nov 10;112(45):E6224–32.

23. Roberts PD. Computational consequences of temporally asymmetric learning rules: I. Differential Hebbian learning. J Comput Neurosci. 1999;

24. Harry Klopf A. A neuronal model of classical conditioning. Psychobiology. 1988;

25. Wörgötter F, Porr B. Temporal Sequence Learning, Prediction, and Control: A Review of Different Models and Their Relation to Biological Mechanisms. Neural Comput. 2005;17(2):245–319.

26. Gluck MA, Parker DB, Reifsnider E. Erratum to: Some biological implications of a differential-Hebbian learning rule. Vol. 17, Psychobiology. 1989. p. 110–110.

27. Turrigiano GG, Leslie KR, Desai NS, Rutherford LC, Nelson SB. Activity-dependent scaling of quantal amplitude in neocortical neurons. Nature. 1998;391(6670):892–6.

28. Van Rossum MCW, Bi GQ, Turrigiano GG. Stable Hebbian learning from spike timing-dependent plasticity. J Neurosci. 2000 Dec 1;20(23):8812–21.

29. Renart A, Song P, Wang X-J. Robust spatial working memory through homeostatic synaptic scaling in heterogeneous cortical networks. Neuron. 2003;38(3):473–85.

30. Pool RR, Mato G. Hebbian Plasticity and Homeostasis in a Model of Hypercolumn of the Visual Cortex. Neural Comput. 2010 Jul 27;1859(7):1837–59.

31. Romo R, Brody CD, Hernández A, Lemus L. Neuronal correlates of parametric working memory in the prefrontal cortex. Nature. 1999 Jun;399(6735):470–3.

32. Machens CK, Romo R, Brody CD. Flexible control of mutual inhibition: A neural model of two-interval discrimination. Science (80-). 2005 Feb 18;307(5712):1121–4.

33. Wu S, Wong KYM, Fung CCA, Mi Y, Zhang W. Continuous Attractor Neural Networks: Candidate of a Canonical Model for Neural Information Representation. F1000Research. 2016 Feb 10;5:156.

34. Wang M, Yang Y, Wang C-J, Gamo NJ, Jin LE, Mazer JA, et al. NMDA Receptors Subserve Persistent Neuronal Firing during Working Memory in Dorsolateral Prefrontal Cortex. Neuron. 2013 Feb;77(4):736–49.

35. Zenke F, Agnes EJ, Gerstner W. Diverse synaptic plasticity mechanisms orchestrated to form and retrieve memories in spiking neural networks. Nat Commun. 2015 Nov 21;6(1):6922.

36. Litwin-Kumar A, Doiron B. Formation and maintenance of neuronal assemblies through synaptic plasticity. Nat Commun. 2014 Dec 14;5(1):5319.

37. Goldman-Rakic P. Cellular basis of working memory. Neuron. 1995 Mar 1;14(3):477–85.

38. Rao SG, Williams G V., Goldman-Rakic PS. Isodirectional Tuning of Adjacent Interneurons and Pyramidal Cells During Working Memory: Evidence for Microcolumnar Organization in PFC. J Neurophysiol. 1999 Apr 1;81(4):1903–16.

39. Constantinidis C, Goldman-Rakic PS. Correlated Discharges Among Putative Pyramidal Neurons and Interneurons in the Primate Prefrontal Cortex. J Neurophysiol. 2002 Dec 1;88(6):3487–97.

40. Strang G. Introduction to Linear Algebra. Wellesley-Cambridge Press; 2017.

41. Rotaru DC, Yoshino H, Lewis DA, Ermentrout GB, Gonzalez-Burgos G. Glutamate Receptor Subtypes Mediating Synaptic Activation of Prefrontal Cortex Neurons: Relevance for Schizophrenia. J Neurosci. 2011 Jan 5;31(1):142–56.

42. Constantinidis C, Franowicz MN, Goldman-Rakic PS. Coding specificity in cortical microcircuits: A multiple-electrode analysis of primate prefrontal cortex. J Neurosci. 2001;21(10):3646–55.

43. Akil AE, Rosenbaum R, Josić K. Synaptic Plasticity in Correlated Balanced Networks. bioRxiv. 2020 Apr 26;

44. Legenstein R, Maass W. Branch-Specific Plasticity Enables Self-Organization of Nonlinear Computation in Single Neurons. J Neurosci. 2011 Jul 27;31(30):10787–802.

45. Ocker GK, Litwin-Kumar A, Doiron B. Self-Organization of Microcircuits in Networks of Spiking Neurons with Plastic Synapses. Latham PE, editor. PLOS Comput Biol. 2015 Aug 20;11(8):e1004458.

46. Mongillo G, Curti E, Romani S, Amit DJ. Learning in realistic networks of spiking neurons and spike-driven plastic synapses. Eur J Neurosci. 2005 Jun;21(11):3143–60.

47. Zenke F, Gerstner W, Ganguli S. The temporal paradox of Hebbian learning and homeostatic plasticity. Vol. 43, Current Opinion in Neurobiology. Elsevier Ltd; 2017. p. 166–76.

48. Chen X, Bialek W. Searching for long time scales without fine tuning. arxiv. 2020 Aug 26;1–13.

49. Machens CK, Romo R, Brody CD. Functional, But Not Anatomical, Separation of “What” and “When” in Prefrontal Cortex. J Neurosci. 2010 Jan 6;30(1):350–60.

50. Murray JD, Bernacchia A, Roy NA, Constantinidis C, Romo R, Wang XJ. Stable population coding for working memory coexists with heterogeneous neural dynamics in prefrontal cortex. Proc Natl Acad Sci U S A. 2017 Jan 10;114(2):394–9.

51. Inagaki HK, Inagaki M, Romani S, Svoboda K. Low-Dimensional and Monotonic Preparatory Activity in Mouse Anterior Lateral Motor Cortex. J Neurosci. 2018 Apr 25;38(17):4163–85.

52. Cueva CJ, Saez A, Marcos E, Genovesio A, Jazayeri M, Romo R, et al. Low-dimensional dynamics for working memory and time encoding. Proc Natl Acad Sci U S A. 2020 Sep 15;117(37):23021– 32.

53. Druckmann S, Chklovskii DB. Neuronal circuits underlying persistent representations despite time varying activity. Curr Biol. 2012;22(22):2095–103.

54. Alemi A, Denève S, Machens CK, Slotine JJ. Learning nonlinear dynamics in efficient, balanced spiking networks using local plasticity rules. In: AAAI Conference. 2018. p. 588–95.

55. Savin C, Triesch J. Emergence of task-dependent representations in working memory circuits. Front Comput Neurosci. 2014;8(MAY):1–12.

56. Hertz J, Krogh A, Palmer RG, Horner H. Introduction to the theory of neural computation. Phys Today. 1991;44(12):70.

57. Chen Z, Haykin S, Eggermont JJ, Becker S. Correlative learning: a basis for brain and adaptive systems. Vol. 49. John Wiley & Sons; 2008.

58. Lim S. Hebbian learning revisited and its inference underlying cognitive function. Curr Opin Behav Sci. 2021;38:96–102.

59. Vogels TP, Sprekeler H, Zenke F, Clopath C, Gerstner W. Inhibitory Plasticity Balances Excitation and Inhibition in Sensory Pathways and Memory Networks. Science (80-). 2011 Dec 16;334(6062):1569–73.

60. Froemke RC. Plasticity of Cortical Excitatory-Inhibitory Balance. Annu Rev Neurosci. 2015;38(1):195–219.

61. Luz Y, Shamir M. Balancing feed-forward excitation and inhibition via hebbian inhibitory synaptic plasticity. PLoS Comput Biol. 2012 Jan;8(1).

